# Point of care influenza testing using Alere-i Influenza A & B assay: a practical assessment

**DOI:** 10.1101/375493

**Authors:** Timothy K Blackmore, James Taylor, Matthew Kelly, Jessica Buckley, Karen Corban, Chor-Ee Tan, Michelle Balm

## Abstract

We assessed the utility of the Alere i Influenza A & B point of care influenza test (Ai-POCIT) with laboratory testing using RT-PCR. 270 adult hospital patients had both Ai-POCIT and laboratory influenza tests conducted on the same sample. Overall, 30% and 32% influenza tests were positive by Ai-POCIT and RT-PCR, respectively. The sensitivity of the Ai-POCIT for influenza A, influenza B and any influenza were 93%, 100%, and 95%, respectively. Specificity was 100% for both viruses, but an 11% test failure rate indicates the need for better training of users. We believe that the use of nasopharyngeal (NP) swabs resulted in the observed high performance of the Ai-POCIT in comparison to other published studies. Ai-POCIT was regarded as very useful by front line clinical staff for clinical decision making and acute bed management.

## Introduction

More than most other respiratory infections, the effective inpatient management of influenza relies upon accurate and rapid laboratory diagnosis so that results can influence clinical decisions on infection control precautions, single room use and antivirals or antibiotic use. This is difficult with laboratory-based testing, especially at weekends and evenings unless the molecular diagnostics laboratory has extended hours of operation and runs the assay more than once a day. Traditional nucleic acid amplification tests (NAAT) are run in batches, further delaying test results that miss the cut-off for being included in the batch. Rapid, one-off NAAT in the laboratory can be helpful (1), but specimen transport to the laboratory and workflow issues may make it difficult to achieve clinically useful turnaround times. It may be logistically more difficult in larger laboratories to identify an urgent swab test for molecular testing from high numbers of other urgent requests.

Driven by requests from clinicians to provide a solution that would better assist with bed management in winter, we prospectively assessed the utility of Alere i Influenza A & B assay, a new point of care influenza test (POCIT) utilising isothermal NAAT. Ai-POCIT was only used on those patients actively considered for hospital admission, and the results were primarily used for room and bed allocation decisions as per infection prevention and control protocols. Testing was performed by selected emergency department (ED) nurses and results were contemporaneously compared with standard laboratory-based RT-PCR testing on the same sample. The testing period included the southern hemisphere winter of 2017 and was conducted in two EDs. The primary aim of the study was to establish whether Ai-POCIT was sufficiently reliable in practice to be introduced as an alternative to laboratory-based influenza diagnostics.

## Materials and Methods

Adult patients being assessed in the ED for admission to medical wards and meeting simple criteria for influenza-like illness (ILI) were included. The criteria for ILI consisted of symptom onset within the past 5 days, with at least one new respiratory symptom (cough, shortness of breath, sore throat, nasal congestion) and at least one new systemic symptom (fevers, rigors, malaise). The study was conducted in winter between July and October 2017, in two ED departments from the largest hospitals in the Wellington region: Wellington and Hutt Hospitals.

A nasopharyngeal (NP) sample was collected using a flocked swab placed into 1 mL of universal transport media (Mini UTM Kit 1 mL, COPAN Diagnostics Inc. Murrieta, CA). The swab was then removed from the tube and tested using Alere i Influenza A & B assay™ (Alere, Waltham, MA, USA). The non-CLIA waived protocol for testing UTM was used, and the remainder of the sample was sent to the laboratory for testing. The amount of material tested in the Ai-POCIT was approximately 75 μL, which is less than the 200 μL recommended by the manufacturer.

The Alere instrument was situated in the ED and was not interfaced with the patient and laboratory information computer systems. Patient labels were not barcoded, requiring patient identification letters and numbers to be entered manually. A limited number of nurses were trained to perform the test, and no in-use training or feedback was provided during the course of the trial.

The laboratory detection of Influenza A and Influenza B was performed by RT-PCR using the RNA Process Control Kit (Roche) for detection of influenza A and B viruses according to WHO/USCDC protocols (CDC Real-time RT PCR Protocol for Detection and Characterization of Influenza, version 2007). The influenza type, cycle threshold value (*C_T_*) and time of result were analysed. The laboratory influenza results were available on the electronic patient management system once checked and released by laboratory staff. Specimens reaching the laboratory by 10 am were tested and reported by 3 pm; those missing the batch were processed on the next working day. Laboratory testing was not routinely available over the weekend.

The results of the Ai-POCIT were immediately available to ED patient care coordinators, and the results were printed and placed in the patient paper record. The difference in turnaround time (TAT) was calculated by comparing the Ai-POCIT result time with the result release time for the laboratory NAAT and was measured in days.

This study was considered to be a laboratory method audit as approved by the Clinical Audit committee of the Capital and Coast and Hutt Valley District Health Boards. The decision to test for influenza remained at the discretion of the treating clinician and there was no change in the method for sample collection. The Alere instruments were provided on loan from the manufacturer, and the manufacturer had no part in the study design, data collection or analysis, or write up. Test kits were paid for utilising departmental research funds. The results were used for bed allocation decisions, but there was no change in policy for the use of antiviral drugs as part of the audit.

## Results

The inclusion criteria requiring patients to be destined for inpatient care meant that the majority of patients had significant comorbidities and reflected the normal range of medical illnesses affecting patients considered for admission to hospital. The study period coincided with the peak period of influenza activity as judged by syndromic surveillance from the Wellington Hospital ED (data not shown).

Three hundred and twenty-one Ai-POCIT results were obtained from 302 patients, of which 270 patients (89%) had laboratory influenza tests conducted on the same sample. Overall, 30% and 32% influenza tests were positive by Ai-POCIT and RT-PCR, respectively. During the same period, 1166 samples were received for laboratory influenza testing. Of patients not having a sample sent to the laboratory, the majority had an Ai-POCIT result of “not detected” (37 of 49, 76%), indicating that the ED staff were less likely to send a sample to the laboratory from a patient for whom they believed influenza had been ruled out. This may have skewed the results assessing the sensitivity of Ai-POCIT, although samples from 163 patients with negative Ai-POCIT results were received by the laboratory.

Thirty-five Ai-POCIT tests from 30 patients failed, leaving 246 results from 243 patients directly comparable. Three patients were repeatedly negative. One patient sample was positive by RT-PCR and negative on initial Ai-POCIT testing, but a repeat Ai-POCIT on the same sample was positive (see below). These results are shown in table 1.

**Table 1.** Concordance of results between Ai-POCIT and laboratory-based RT-PCR for the 247 patients who had both tests performed. A = influenza A detected, B = influenza B detected, AB = influenza A and B detected, ND = not detected

| Ai-POCIT Result | Laboratory RT-PCR Result |  |  |  | Total |
| --- | --- | --- | --- | --- | --- |
|  | A | A + B | B | ND |  |
| A | 50 |  |  |  | 50 |
| B |  | 1* | 23 |  | 24 |
| ND | 4 |  |  | 168 | 172 |
| Total | 54 | 1 | 23 | 168 | 246 |

The sensitivity of the Ai-POCIT for influenza A, influenza B and any influenza were 93%, 100%, and 95%, respectively, compared to the laboratory RT-PCR assay. There were no false positives for either virus. One mixed, predominately influenza B infection was detected as B only by Ai-POCIT.

Three of the four false negative influenza A results by Ai-POCIT were seen in those with lower viral copies as judged by *C_T_* values on RT-PCR testing (fig. 1). The false negative result with a *C_T_* value of 28 was repeated by a different operator on the same sample, and was found to be positive, indicating an unrecognised process error on the first Ai-POCIT test.

**FIGURE 1.**
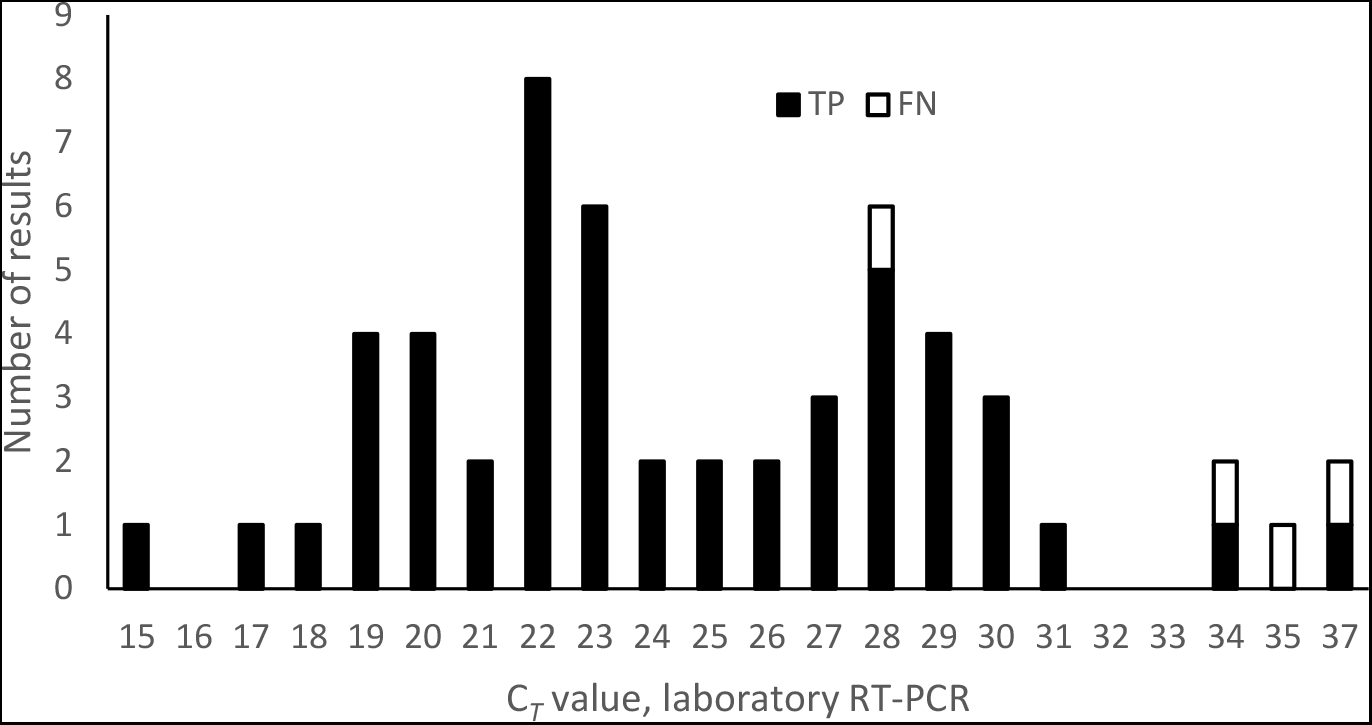
Frequency distribution of cycle threshold (*C_T_*) values for samples positive for influenza A virus in the laboratory RT-PCR test. *C_T_* values were rounded to the nearest whole number. True-positive (TP) and false-negative (FN) Ai-POCIT test

There were 35 tests that failed on Ai-POCIT (11%). There were 29 trained users of the device, but 60% of failed results were performed by three users. Process review suggested that the predominant sources of procedural error were from exceeding the sample receiver warm-up time, and incomplete sample transfer. There was no correlation between the number of tests performed by each user and the number of errors. In addition to test failures, 3% of samples had inconsistencies in sample identification, none of which were sufficient to prevent matching with laboratory results.

As expected, the turnaround time for Ai-POCIT was much shorter than for laboratory-based testing. Only 23% and 51% of laboratory results overall were available the same day or next day, respectively. The timeliness of the laboratory results depended on the day and time the specimen was sent to the laboratory, as shown in figure 2.

**FIGURE 2.**
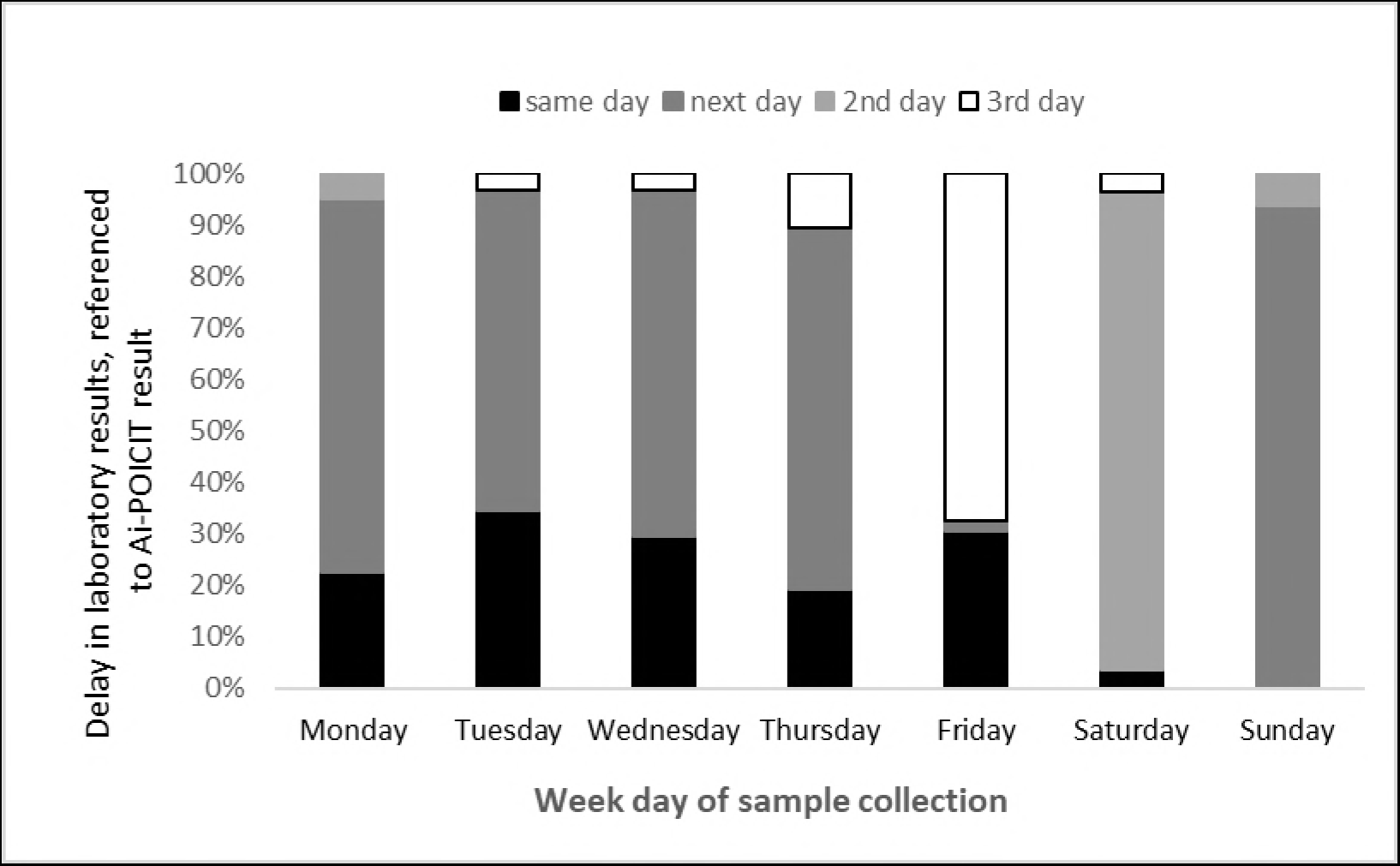
Availability of laboratory influenza test results relative to Ai-POCIT tests results according to the day of the week that Ai-POCIT was performed.

Of the 302 patients, 203 would have received an influenza swab previously, and then required droplet isolation.

The proportion of patients receiving laboratory results by day (figure 2) show that 72 after 1 day, 18 after 2 days and 16 after 3 days, respectively, would have been removed from contact precautions if hospital infection prevention and control guidelines were followed. This is a total of 156 isolation bed days. If all patients meeting criteria for ILI requiring admission were tested and isolated pending results, 233 isolation bed days would have been saved.

If laboratory PCR was performed daily, the total isolation bed days would be 95, calculated by proportion of weekday samples receiving next day laboratory results.

Treatment was not a focus of our study, but 171 patients received antibiotics, predominately amoxicillin and azithromycin, consistent with local community-acquired pneumonia guidelines.

## Discussion

Testing for influenza virus may be performed for a variety of clinical or epidemiological reasons, but we recommend testing in our hospitals only for bed allocation decisions for and treatment of early presentation of ILI. It is common for “winter bed block” to occur whereby patient admission to isolation rooms or cohort areas can have a major impact on patient flow and emergency department waiting times. Prior to conducting this audit, influenza RT-PCR had been available only 5 days a week. The testing was conducted in batches, with a sample cut-off time of 10 am, with results available around 2 pm. This required patients with ILI to be placed in isolation pre-emptively, often for prolonged periods: there was a strong desire by clinicians and administrators to improve diagnostic turnaround times.

The availability of nucleic acid amplification-based POCIT offered a potential solution, because of the claimed improved sensitivity over other rapid methods (2). Due to staffing and workflow issues, it would not have been possible to provide laboratory-based test results significantly more quickly without prohibitively increasing costs. This audit was therefore conducted to examine the practical issues involved in POCIT and to validate the Ai-POCIT in our clinical setting.

Our results indicate that the Ai-POCIT test provides prompt, accurate results and the ED staff were happy to conduct the test themselves instead of sending a sample to the laboratory. Our sensitivity and specificity results are comparable to a recent meta-analysis, with better performance of Ai-POCIT against influenza B than influenza A (2), and like others we found that the majority of false negative influenza A results occurred at higher *C_T_* values by RT-PCR (3). It is tempting to speculate that such patients may be less infectious to others because of lower viral loads in respiratory secretions and therefore of less concern if missed on testing. There have been two research publications regarding Ai-POCIT since the meta-analysis by Merkx et al. In a multicentre study using throat swabs, Davis et al found only a 77% sensitivity compared to RT-PCR, but they did not distinguish between influenza types. There was also a large variation in test performance between sites (4). Young et al found Ai-POCIT less sensitive than the Cobas Liat POCIT, using stored UTM samples (5). There has also been an excellent review of rapid tests for influenza recently published (6).

An advantage of our study design was that each sample was tested by both RT-PCR and Ai-POCIT, allowing a direct comparison of test performance. Other published studies of Ai-POCIT have used a variety of methods regarding sample type, split or separate samples, sample storage and relative timing of alternative tests (3, 5, 7–10).

Testing the same sample in parallel meant that the samples were in effect split, and that the Ai-POCIT was run on the non-CLIA-waived protocol used for testing aspirates. We would expect better sensitivity for both assays if all material collected on the NP swab was placed into either the receiving chamber (Ai-POCIT) or extraction (RT-PCT), but this may not be significant as suggested by the higher observed sensitivity in this compared with other studies (2, 6). Similarly, RT-PCR was conducted on UTM in which the swab was twirled and then removed: a process which may have released less target material into the media. The laboratory normally receives NP swabs in 1mL of UTM, which is then vortexed before using 200 μL for RNA extraction.

It is likely that by utilising NP swabs we maximised sensitivity of both assays. The impact of lower sensitivity of Ai-POCIT compared to RT-PCR would be reduced by collecting the best sample with the most virus present. Frazee et al showed that the best sample is the NP swab, but the best balance between comfort and sample quality is found with midturbinate swabs (10).

This study provides detail about errors occurring during a season of Ai-POCIT use. The 11% failure rate significantly adds to costs and shows the importance of feedback and retraining of staff. The data log from the instrument makes it easy to provide individual staff feedback which would be important to maintain quality. The first timed step of warm-up prior to introduction of the sample is 5-10 minutes, during which other jobs may be started. Time-outs occur as a result of the staff member becoming distracted. Anecdotally, this error appeared to be more frequent when ED staff were conducting a test for another clinical area, rather than their own patient. They found that it was easier to become distracted during the warm up phase. The use of a timer on a cellular phone helps to reduce these errors. Another common mistake of not “clicking” the transfer device properly and failing to observe the orange indicator tab move is something that can be minimised with good training. Finally, the clerical errors observed would be completely removed with the use of barcoded samples. All of these errors appear to be largely resolved in the subsequent season of use, through better training.

This audit is one of the largest published of POCIT to date, and the results are consistent with other published studies (3, 7). We experienced a slightly higher activity influenza season than usual, as judged by syndromic surveillance in the Wellington Hospital ED. The 30% positivity rate in adult patients admitted to medical wards indicates that the loose definition of ILI employed in this study was reasonably effective: the positive rate from routine RT-PCR was similar overall, using normal testing practice. The predominant circulating viruses in New Zealand at the time were A(H3N2), B/Yamagata and B/Victoria lineage (11). It may be that POCIT performance would be affected by variation of circulating strains, and so it will be important to continue to monitor performance regularly (12).

It can be argued that other respiratory viruses should be tested for, with some advocating that diagnosing a viral respiratory tract infection may useful for antibiotic stewardship. Apart from the lack of availability of rapid multiplex respiratory PCR panels, we would not be able to provide bespoke infection prevention measures for all respiratory viruses. We currently do not routinely perform multiplex PCR routinely on patients with respiratory tract infection, with the exception of those admitted to haematology-oncology. The introduction of Ai-POCIT will allow influenza testing to be performed expeditiously, and for our laboratory to only perform multiplex respiratory pathogen testing.

POCIT may be the best way to achieve rapid, reliable testing for influenza in larger facilities, but this may not be cost effective for smaller facilities where rapid laboratory-based RT-PCR tests may be more cost effective and easier to manage (8).

The saving of 156 isolation bed days had a noticeable effect on hospital bed flow, because of increased availability of isolation facilities. This in turn allowed better patient flow through the ED with beneficial effects for the whole hospital. Early exclusion of influenza also reduced the number of patients in contact precautions, potentially improving patient safety(13).

Even a shift to a daily laboratory PCR would not have provided this level of improvement, as 95 isolation bed days would still be required due to the once daily nature of testing.

The ready availability of Ai-POCIT in the ED ensured that testing happened as intended. In particular the senior nurses performing the tests had an incentive to do so, as without the test many patients would have required a wait for an isolation room. Point of care testing should be delivered at the point of care, not merely using this technology as a faster and easier method of running a laboratory. Finally by having a cost of testing (of clinician time), this will have had an automatic demand management effect. It is only utilised when appropriate, and not for patients who would be discharged or otherwise do not need a precise microbiological diagnosis.

Only 18 patients received oseltamivir, 3 of whom were Ai-POCIT negative. Two of these patients were also negative on laboratory influenza testing, and it is unclear why this was prescribed. The final patient prescribed oseltamivir was immunosuppressed. This was prescribed following a positive laboratory PCR despite a negative Ai-POCIT.

We have shown that POCIT has dramatically improved the utility of influenza testing in the hospital setting, with major effects on bed flow and ED department waiting times. The availability of sensitive POCIT such as Ai-POCIT is also likely to change the way care is delivered, with a realistic option of treating confirmed influenza quickly with antiviral medication (14). Few hospitalised patients in the past were diagnosed with influenza early enough to benefit from antivirals. Our data show that over 75% of patients should now get their results one to three days earlier, dramatically improving chances for useful clinical intervention. This study indicates that Ai-POCIT provides reliable results when NP swabs are tested; we therefore now recommend treatment in patients hospitalised with influenza requiring hospitalisation within 3 days of symptom onset.

## Acknowledgements

This study was funded from Infection Services departmental research funds, Capital and Coast District Health Board.

TB has received a speakers honorarium from Alere/Abbott diagnostics. No other financial disclosures

